# Regulation of mitochondrial DNA homeostasis by a mitochondrial microprotein

**DOI:** 10.64898/2026.02.20.706844

**Authors:** Ikram Ajala, Damien Lipuma, Grégoire Bonnamour, Benoît Vanderperre

## Abstract

The actin cytoskeleton and non-muscle myosins coordinate a wide array of cellular functions, yet their contribution to mitochondrial genome maintenance remains poorly understood. Here, we identify AltSLC35A4, a conserved protein encoded by an alternative open reading frame (AltORF), as a novel regulator of mitochondrial DNA (mtDNA) homeostasis. Using co-immunoprecipitation and mass spectrometry, we show that AltSLC35A4 interacts with actomyosin cytoskeletal regulators including MYH9 and MYH10, which have been previously implicated in mtDNA regulation. Loss of AltSLC35A4 increases mtDNA copy number and causes a dispersed spatial distribution of mitochondrial nucleoids, leading to the accumulation of extramitochondrial DNA puncta. Theses alterations occur without detectable changes in TFAM expression and mitochondrial membrane potential, suggesting that the mtDNA dysregulation is independent of mtDNA replication and transcription and mitochondrial bioenergetic state. Our findings uncover a previously unrecognized role for AltSLC35A4 in mitochondrial nucleoids regulation and highlight the functional importance of altORF-encoded proteins in mitochondrial biology.

## Introduction

The actin cytoskeleton orchestrates a wide range of essential cellular functions, including intracellular trafficking, membrane remodeling, and organelle positioning^1^. These dynamic processes rely on the polymerization and turnover of actin filaments, tightly controlled by actin-binding proteins (ABPs) that regulate filament nucleation, elongation, severing, and disassembly ^1^. Recent studies have expanded the scope of actin cytoskeleton’s role beyond the cortex and cytoplasm, highlighting its involvement in mitochondrial dynamics, mitochondrial biogenesis, motility, and quality control ^2,3^.

Non-muscle myosins are major contributors to actin-driven mechanical activity. These ATP-dependent motor proteins interact with filamentous actin to produce contractile tension and facilitate intracellular transport^1^. In particular, the class II myosins MYH9 and MYH10 (non-muscle myosin heavy chain IIA and B, respectively) contribute to cellular processes such as cytokinesis, vesicle movement, adhesion dynamics, and mechanical signal transduction^4^. Both have been implicated in maintaining cytoskeletal architecture and coordinating cellular responses to stress^4^. Importantly, several studies have revealed that myosins and actin can regulate mitochondrial DNA (mtDNA) homeostasis through both direct and indirect mechanisms. Cytosolic myosin proteins influence mitochondrial morphology and fission dynamics by coordinating actin polymerization at the outer membrane, which in turn affects nucleoid positioning and segregation during organelle division^5,6^. Recent findings have extended the influence of the actomyosin cytoskeleton beyond the mitochondrial surface to the organelle interior. Super-resolution microscopy has revealed β-actin–containing filaments within the mitochondrial matrix, where they participate in the maintenance of mtDNA integrity and nucleoid organization^7^. Loss of β-actin results in altered mtDNA copy number, disordered nucleoid distribution, and increased vulnerability to mitochondrial stress, highlighting a direct role for actin in mtDNA transcription and quality control ^7^. Moreover, myosin II has been detected in association with purified mitochondrial nucleoids, and its silencing produces mtDNA abnormalities. Together, these observations support the existence of an internal “mitoskeleton” composed of actin and myosin elements that contribute to mtDNA segregation and inheritance, providing an additional level of spatial organization and regulation within mitochondria^1^. While the involvement of actin and myosin in shaping mitochondrial morphology and dynamics is increasingly recognized ^3,5,8^, the mechanisms linking these cytoskeletal components to the regulation of mitochondrial DNA (mtDNA) remain largely unexplored. This is especially true at the level of the mitochondrial inner membrane (IMM), where regulatory factors for mtDNA homeostasis are still being identified.

MtDNA is packaged into nucleoids whose structural organization and spatial distribution both safeguard genome stability and ensure proper mitochondrial function in mammalian cells. Super-resolution imaging established that human nucleoids are compact structures that often harbor a single mtDNA molecule, emphasizing how their spacing and maintenance are tightly regulated in healthy cells^9^. Disruption of these processes undermines oxidative phosphorylation and can precipitate cellular stress phenotypes^10^.

MtDNA maintenance depends on the balance between replication and turnover as well as on nucleoid-membrane coupling. The transcription factor TFAM is a key determinant of mtDNA copy number *in vivo*, while IMM factors such as ATAD3A help organize nucleoid positioning and trafficking, linking genome maintenance to mitochondrial ultrastructure^11^.

Beyond canonical gene annotations, a growing alternative proteome, comprising proteins translated from alternative or small open reading frames (AltORFs/sORFs) is emerging as a source of previously unrecognized regulators, including mitochondrial proteins^12^. Recent functional studies underscore that alternative proteins can localize to mitochondria and modulate the physiology of this organelle, suggesting an underappreciated layer of mitochondrial regulation ^13,14^.

Us and others have uncovered AltSLC35A4, a previously uncharacterized protein encoded by an alternative open reading frame (altORF), as a protein of the IMM^15,16^. Despite its evolutionary conservation and distinct mitochondrial localization, the functional role of AltSLC35A4 has remained elusive. Our prior work demonstrated that AltSLC35A4 deficiency renders cells more susceptible to oxidative stress^15^. Recent studies have confirmed that the loss of AltSLC35A4 (a.k.a. SLC35A4-MP for SLC35A4 microprotein) in human cells reduces basal, maximal and ATP-linked oxygen consumption rates, while its overexpression enhances mitochondrial membrane potential and respiratory capacity, supporting a key role in oxidative metabolism^16^. At the organismal level, brown adipose tissue of AltSLC35A4 knock-out (KO) mice display altered mitochondrial lipid composition, decreased cardiolipin and phosphatidylethanolamine content, and impaired mitochondrial respiration, accompanied by inflammatory remodeling under metabolic stress ^17^.

In the present study and through co-immunoprecipitation, transcriptomic profiling, and high-resolution imaging, we reveal that AltSLC35A4 engages with key cytoskeletal regulators, including MYH9 and MYH10 in human 143B cells. Loss of AltSLC35A4 perturbs the expression of actomyosin-related genes and inflammatory pathways, elevates mitochondrial DNA copy number, and disrupts the spatial organization of mitochondrial nucleoids. Notably, cells lacking AltSLC35A4 accumulate extranuclear mtDNA puncta, a phenotype reversed by reintroducing the protein. These findings suggest that AltSLC35A4 may functionally coordinate cytoskeletal dynamics and mitochondrial nucleoid organization, thereby contributing to mitochondrial genome maintenance. Altogether, our results identify AltSLC35A4 as a novel regulator at the interface between the cytoskeleton and the IMM, advancing our understanding of how myosin-dependent mechanisms may influence mitochondrial genome maintenance.

## Experimental procedures

### Cell culture

143B and HEK293T cells were cultured in Dulbecco’s Modified Eagle Medium (DMEM) (Wisent, #319-005-CL) containing 10% fetal bovine serum (FBS) (Wisent, #080-450) and 1% penicillin-streptomycin (Bio Basic, #PB0135,-# SB0494) under standard conditions (37°C in a humidified atmosphere with 5% CO2). Cells were routinely tested for mycoplasma contamination.

### Coimmunoprecipitation-Mass spectrometry

For co-immunoprecipitation of AltSLC35A4-3xHA, cells stably expressing AltSLC35A4-3xHA were cultured in 150 mm plates until they reached 80% confluency. The TNI buffer was prepared by combining 50 mM Tris (pH 7.5), 0.5% Igepal, 250 mM NaCl, 1 mM EDTA, along with 0.3 µM Aprotinin (Cederlane, #14716-10), 4.2 µM Leupeptin (Cederlane, #LEU001.10), and a broad-spectrum phosphatase inhibitor cocktail (Boster Immunoleader, #AR1183). Cells were rinsed thoroughly with at least 10 mL of 1× PBS, then lysed with 1.5 mL of ice-cold TNI buffer per culture dish. After a 20-minute incubation on ice with gentle agitation on an orbital shaker, cells were scraped and the lysate was transferred to a 1.5 mL tube and centrifuged at 18,000 × g for 30 minutes at 4°C to obtain the supernatant. The resulting clarified supernatant was then collected, and a 50 µL aliquot was saved as input fraction. Supernatant was transferred to prewashed HA-magnetic beads (ThermoFisher Scientific, #88837) and incubated for 2 hours at 4°C. Beads were collected using a magnetic rack and 50 µL aliquot of the unbound fraction was kept. Beads were transferred to a new microcentrifuge tube and washed three times with TNI lysis buffer, followed by two washes with TN (without Igepal) lysis buffer. A portion of the final wash was preserved for Western blot and silver stain validation, with the remaining beads sent for mass spectrometry analysis at the Proteomics and Molecular Analysis Platform of McGill University Health Center - Research institute (RI-MUHC).

### Western blot, immunofluorescence, confocal and super resolution microscopy

Western blot, immunofluorescence and confocal microscopy analyses were carried out as previously described^15^. Live cell imaging was performed under controlled conditions as detailed in the JC-1 assay section.

STimulated Emission Depletion (STED) super-resolution imaging was performed on a Leica Stellaris 8 STED microscope equipped with a tunable White Light Laser (440–790 nm) for excitation and a pulsed 775 nm STED depletion laser. Images were acquired using a 100X oil immersion objective (NA 1.4) at room temperature. Pixel size ranged from 12 to 18 nm, achieved through a digital zoom between 2 and 6x. Fluorescence was detected using hybrid detectors (HyD) with spectral windows optimized for the fluorophores used. Acquisition parameters included a dwell time of 700 ns per pixel and line averaging of 10.

Antibodies used are listed in Supplementary Table I.

### RNA extraction and RNA-Seq analysis

A total of 0.1 × 10 cells were seeded per well in 6-well plates in triplicate until they reached 80% confluency. Total RNA was extracted using the miRNeasy Mini Kit (Qiagen, Cat# 217004) according to the manufacturer’s instructions. RNA samples were then sent to the Genomics Platform of CERMO-FC for quality control assessment and mRNA enrichment, followed by RNA-Seq libraries preparation. Sequencing of the RNA-seq libraries (25 million paired-end reads per sample, 2 × 100 bp) was subsequently performed at Génome Québec using the Illumina NovaSeq 6000 (kit S1, v1.5). Four RNA samples were used for RNA sequencing, including two biological replicates from wild-type (WT) cells and two from AltSLC35A4 KO cells. Raw sequencing data were processed and analyzed using the Galaxy web-based platform (http://usegalaxy.org). Quality trimming of raw reads was performed using Trimmomatic (version 0.39), and read quality was assessed using FastQC (version 0.74). Alignment to the human reference genome (GRCh38) was carried out using HISAT2 (version 2.2.1). Gene-level quantification was performed with FeatureCounts (version 2.0.8), followed by downstream analysis using annotatemyID (version 3.18.0) and limma (version 3.58.1). Differentially expressed genes (DEGs) were identified based on a log fold change ≥ 1 or ≤ −1 and an adjusted p-value (FDR) < 0.01. Final DEG lists were treated using Microsoft Excel. Gene Ontology enrichment and network analyses were carried out using the STRING database (version 12.0), which integrates known and predicted protein - protein interactions. DEGs were mapped to STRING to assess Reactome pathways. Networks were visualized based on false discovery rate (FDR)-adjusted p-values.

### qPCR

Cells were cultured in a 24-well plate until reaching 80% confluency. Total DNA was extracted using the Quick Extract Kit (Cederlane, #QE09050) according to the manufacturer’s instructions. Primers targeting the mitochondrial *COX2* and nuclear *ACTB* genes were designed (*b-actin-F*: 5′-CTTCCCTCCTCAGATCATTG-3′; *b-actin-R*: 5′-TTGTCAAGAAAGGGTGTAAC-3′. *COX2-F*: CCTACATACTTCCCCCATTA; *COX2-R:* CAGATTTCAGAGCATTGACC). qPCR was conducted using a Bio-Rad qPCR system (CFX96^TM^) with primers at a final concentration of 0.25 µM and Luna qPCR Master Mix (NEB, #M3003L) according to the manufacturer’s instructions. All samples, including nuclease-free water as a no-template control, were run in technical triplicate at a DNA concentration of 5 ng per replicate. After normalization, the CT values for the mitochondrial gene target were analyzed using the 2^-ΔΔCt^ method to assess gene expression and calculate the relative change in mitochondrial DNA copy number. Specifically, ΔCt was determined by subtracting the CT value of the *ACTB* gene from that of the *COX2* gene (ΔCt = Ct^mito^ - Ct^nuc^). Subsequently, ΔΔCt was calculated by subtracting the ΔCt of the sample from the ΔCt of the WT control, allowing for the calculation of relative change in mitochondrial DNA copy numbers based on ΔΔCt (N = 2^-ΔΔCt^) to estimate the ratio of mitochondrial to nuclear DNA.

### Mitochondrial membrane potential assay using JC-1

Twenty-four hours prior to the experiment, 10,000 cells were seeded in each well of a µ-Slide 8 well ^high^ Glass Bottom (ibidi - cells in focus, #80807). After 24 hours, cells were washed once with PBS and stained with JC-1 dye (5,5′,6,6′-tetrachloro-1,1′,3,3′-tetraethylbenzimidazolcarbocyanine iodide; final concentration: 7.6 μM in DMSO) (StressMarq Biosciences, SIH-562-5MG) for 15 minutes at 37 °C. Where indicated, cells were pretreated with 5 μM carbonyl cyanide-p-trifluoromethoxyphenylhydrazone (FCCP) (Sapphire Bioscience, #A16545-5) for 5 minutes prior to JC-1 labeling. Live-cell imaging was performed using Nikon A1 confocal microscopy, with cells maintained in a micro-incubator under controlled conditions (temperature 37°C and CO 5%). Images were acquired using Plan Apo λ 60× oil-immersion objectives (NA 1.4; refractive index 1.515) in a single focal plane. JC-1 fluorescence was recorded using a 488 nm laser (emission collected at 525/50 nm) for the monomeric green form and a 561 nm laser (emission collected at 593/46 nm) for the red J-aggregate form. The mitochondrial membrane potential (Δψm) was quantified by calculating the red-to-green fluorescence intensity ratio.

### Mitochondrial nucleoids analysis with CellProfiler

Confocal Z-stacks (7 optical sections, 1.34 µm step size, total depth ≈ 8 µm) were acquired on a Nikon A1 confocal microscope equipped with a Plan Apo λ 60× oil immersion objective (NA 1.4, refractive index 1.515). Maximum intensity projection images of confocal Z-stacks stained for dsDNA (mtDNA), mitochondria (TOM20), and nuclei (Hoechst 33342) were analyzed using CellProfiler (version 4.2.8, Broad Institute). Images were first pre-processed in Fiji to generate maximum intensity projections and imported into CellProfiler. The analysis pipeline included enhancement of punctate structures using the EnhanceOrSuppressFeatures module (Enhance Speckles) to emphasize nucleoids, followed by segmentation of individual nucleoids with IdentifyPrimaryObjects (adaptative threshold strategy with minimum cross-entropy as thresholding, declumping based on intensity). Nuclei were detected in the Hoechst channel, and cell boundaries were defined as secondary objects using the mitochondrial channel and the Propagation method. The mitochondrial DNA signal overlapping with nuclei was masked out to exclude nuclear regions and the remaining nucleoids were then related to their parent cell using RelateObjects. In addition, extracellular nucleoids, defined as mtDNA puncta detected outside the cell boundaries, were quantified separately to assess mitochondrial DNA release events.

Quantitative features such as nucleoids count per cell, mean nucleoid area, and mean nucleoid intensity were extracted with MeasureObjectSizeShape and MeasureObjectIntensity, while cell area was determined using MeasureImageAreaOccupied. Data were exported to spreadsheets for further analysis in Excel. For each field, the mean number of nucleoids per cell and nucleoid density (number of nucleoids per µm² of cell area) were calculated. All segmentation parameters (object diameters, thresholds, and feature size) were optimized on a representative training set and applied uniformly to all images.

### Data availability and statistics

All data supporting the findings of this study are available from the corresponding author upon reasonable request. The custom CellProfiler pipelines used for image quantification and analysis can also be provided upon request. All statistical analyses, except for enrichment analysis of RNA-Seq and mass spectrometry data, were performed with GraphPad PRISM (version 10.4.1) using one-way ANOVA with adequate multiple comparisons tests (specified in figures legends) to assess differences between groups. Data are presented as mean ± standard deviation (SD) and statistical significance was defined as p < 0.05 unless otherwise indicated.

## Results

### AltSLC35A4 interacts with myosin heavy chains MYH9 and MYH10

To identify potential interactors of AltSLC35A4 and gain insights into its biological function, we performed co-immunoprecipitation (coIP) followed by mass spectrometry using 143B cells expressing AltSLC35A4 fused to an N-terminal 3xHA tag. We observed a significant enrichment of cytoskeletal proteins compared to control cells, including several actin-associated myosins such as MYH9, MYH10, MYO1B, and MYO1C, as well as actin-binding components (ACTBL2, TPM3, TPM4, MYL6, and MYL12A) (Fig. 1A). Gene Ontology enrichment analysis of the AltSLC35A4 interactome further supported the association of this alternative protein with actin-related processes, highlighting GO terms such as actin filament organization, cytoskeleton organization, and supramolecular fiber assembly (Fig. 1B). We confirmed the interaction of AltSLC35A4 with MYH9 and MYH10 by coIP and Western blot (Fig. 1C–D). Consistent with these findings, immunofluorescence microscopy showed partial colocalization of 3xHA-AltSLC35A4 with MYH9 (Fig. 1E). Together, these results suggest that AltSLC35A4 physically associates with the actomyosin cytoskeleton, presumably in the mitochondrial interior given that AltSLC35A4 localizes to the IMM, potentially linking mitochondrial structure to actin-dependent organization and dynamics.

**Figure 1:**
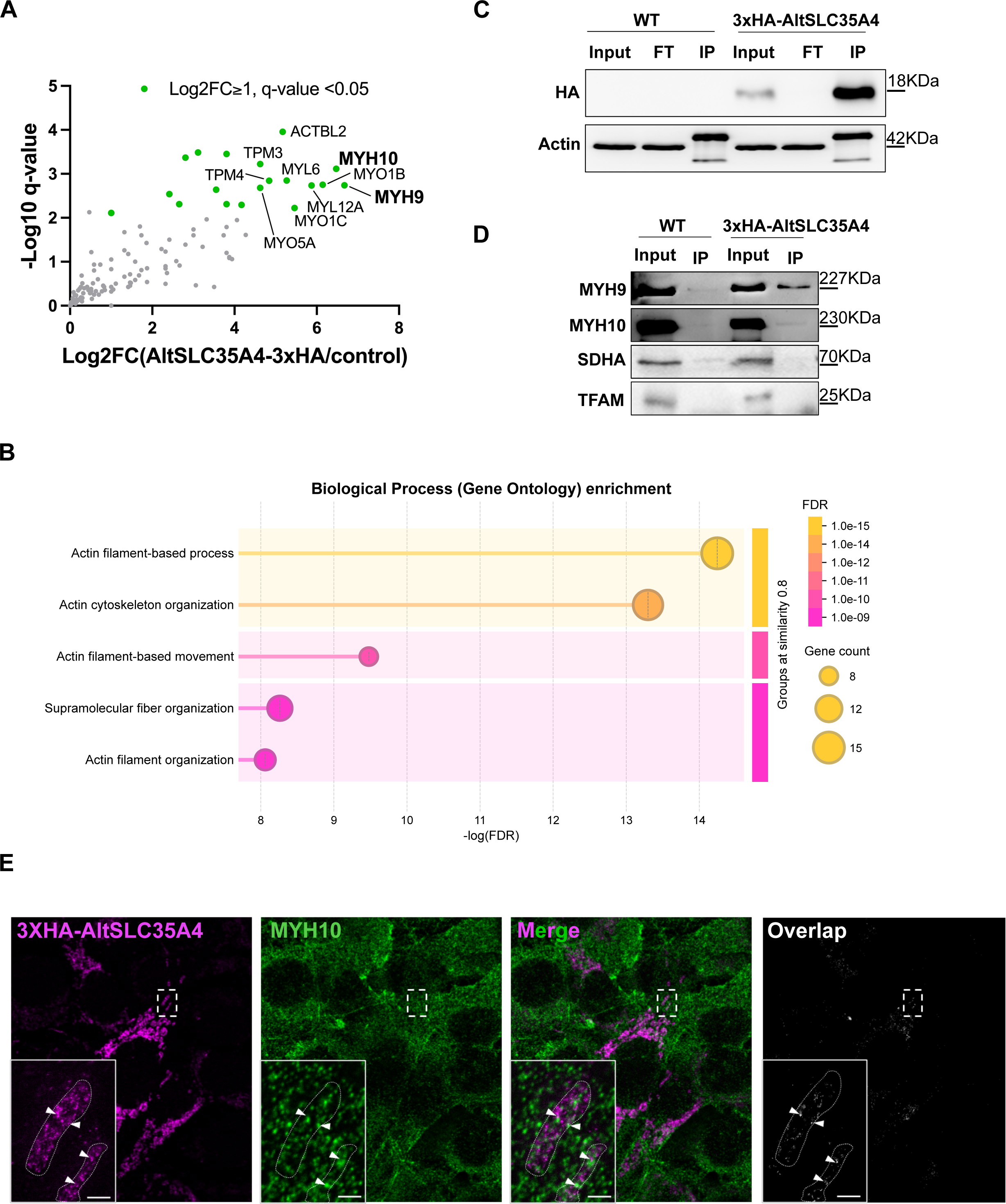
AltSLC35A4 interacts with actomyosin cytoskeletal proteins. (A) Volcano plot of proteins enriched in AltSLC35A4-3xHA immunoprecipitates compared to control cells, as determined by LC-MS/MS. Significantly enriched interactors (Log2FC ≥ 1, q-value < 0.05) are shown in green, with major cytoskeletal proteins (MYH9 and MYH10) highlighted. (B) Gene Ontology (GO) enrichment analysis of significantly enriched proteins revealed strong overrepresentation of actin cytoskeleton–related biological processes, including actin filament organization and actin-based movement. (C) Co-immunoprecipitation of AltSLC35A4-3xHA confirmed successful enrichment of the tagged protein compared to wild-type (WT) cells, with actin used as loading control. FT, flow-through; IP, immunoprecipitate. (D) Immunoblot validation of AltSLC35A4-3xHA interactions with non-muscle myosins MYH9 and MYH10, with SDHA and TFAM serving as negative control for pulldown of mitochondrial material. (E) Super resolution microscopy showing partial colocalization between AltSLC35A4-3xHA (magenta) and MYH9 (green). The merged image shows discrete regions where both signals overlap (shown with arrow heads). The Overlap panel displays only the pixels positive for both markers, highlighting areas of true colocalization. Insets show magnified views of the boxed regions. Scale bar: 1 µm.

### Disruption of AltSLC35A4 leads to mtDNA accumulation and leakage

The identification of cytoskeletal proteins (MYH9 and MYH10) as AltSLC35A4 interactors suggested that this microprotein could be involved in intramitochondrial mechanisms involving actomyosin dynamics. Interestingly, an intramitochondrial pool of these cytoskeletal proteins exists and is thought to contribute to the maintenance of mtDNA nucleoids^18^. In addition, AltSLC35A4’s localizes within the IMM and several IMM-resident proteins have been reported to regulate mtDNA. Thus, we reasoned that AltSLC35A4 might contribute to mtDNA maintenance. We therefore assessed mtDNA content and spatial organization in AltSLC35A4 knockout (AltSLC35A4-KO) cells. Quantitative PCR analysis revealed a significant increase in mtDNA copy number in AltSLC35A4-KO cells compared to wild-type (WT) cells, which was restored upon stable re-expression of AltSLC35A4 (AltSLC35A4-KO resc) (Fig. 2A), suggesting a role for this protein in mtDNA homeostasis. Confocal imaging of mitochondria and mtDNA nucleoids further reinforced this phenotype, revealing a higher density of mtDNA foci in KO cells together with prominent extramitochondrial DNA puncta (Fig. 2B). High-magnification views confirmed that these puncta reside in the cytosol adjacent to mitochondria (Fig. 2B). Automated quantification using a custom CellProfiler pipeline demonstrated a significant increase in the number of nucleoids per cell in KO cells (Fig. 2C), while nucleoid size remained unchanged across genotypes (Fig. 2D). Consistent with these observations, KO cells also exhibited an increase in extramitochondrial DNA puncta, which was rescued upon AltSLC35A4 re-expression (Fig. 2E), whereas puncta size did not differ between genotypes (Fig. 2F).

**Figure 2:**
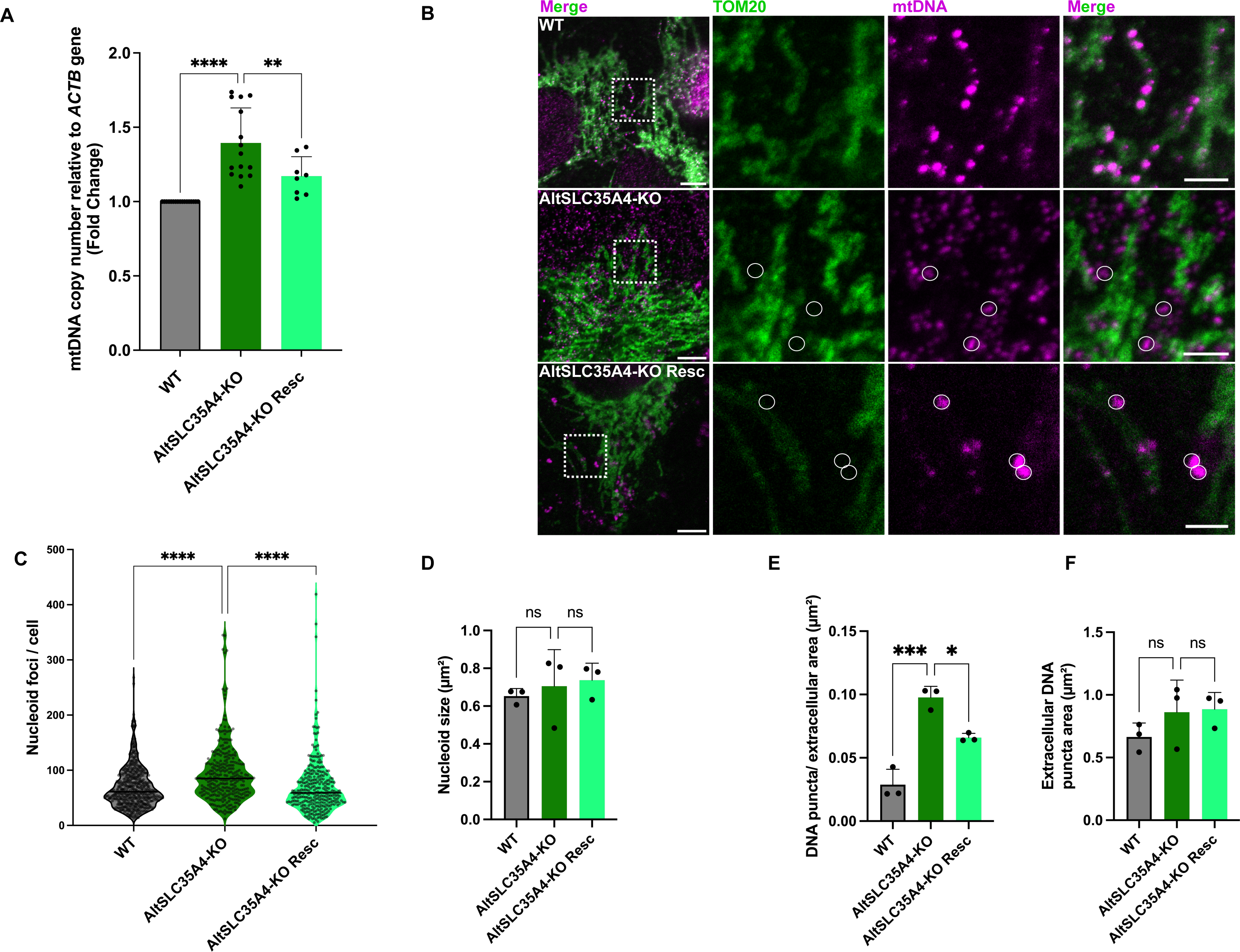
AltSLC35A4 deficiency increases mtDNA copy number and induces extramitochondrial and extracellular DNA puncta. (A) Quantification of mitochondrial DNA (mtDNA) copy number by qPCR using *ACTB* as a nuclear reference gene. AltSLC35A4-KO cells display a significant increase in mtDNA copy number compared to wild-type (WT) cells, which is rescued by AltSLC35A4 re-expression. (B) Confocal microscopy of WT, AltSLC35A4-KO, and rescue cells stained for TOM20 (green), mtDNA (magenta, dsDNA), and nuclei (blue, Hoechst). AltSLC35A4-KO cells show an increase in nucleoids number and extramitochondrial DNA puncta. Scale bar: 10 µm. (C) Quantification of mitochondrial nucleoids per cell using CellProfiler analysis shows a significant increase in AltSLC35A4-KO cells, rescued by AltSLC35A4 expression. (D) Nucleoid size shows no significant difference among genotypes (E) Extramitochondrial DNA puncta area normalized to total extracellular area. (F) Mean area of individual extracellular DNA puncta is unchanged. Data represent n=3 biological replicates. For image-based quantifications, between 77 and 202 cells were analyzed per replicate. Values are presented as mean ± SD. Statistical significance was determined by one-way ANOVA with Tukey’s multiple comparison test; *, p < 0.05; **, p < 0.01; ****, p < 0.0001; ns, not significant.

Together, these findings demonstrate that loss of AltSLC35A4 leads to mitochondrial DNA accumulation and mislocalization into extramitochondrial and extracellular compartments, suggesting that this protein is required to control mitochondrial genome copy number and prevent mtDNA release into the cytosol and ultimately in the extracellular space.

### AltSLC35A4 deficiency alters mtDNA organization independently of TFAM expression or mitochondrial mass

Because the increase in mtDNA copy number and extramitochondrial DNA puncta observed in AltSLC35A4-KO cells could result from changes in nucleoid packaging or TFAM abundance^19,20^, we next assessed TFAM protein levels and distribution. Western blot analysis showed that total TFAM expression remained unchanged across WT, AltSLC35A4-KO, and rescue cells (Fig. 3A-B), indicating that the observed mtDNA phenotype is not due to altered TFAM protein abundance. Confocal microscopy of TOM20 and TFAM co-staining revealed no overt differences in TFAM distribution between conditions (Fig. 3C). Quantitative image analysis further confirmed that both total and mean TFAM fluorescence intensity, normalized to mitochondrial area, were comparable among WT, AltSLC35A4-KO, and rescue cells (Fig. 3D, E). To determine whether the observed mtDNA increase could instead reflect an expansion of the mitochondrial network, we next assessed mitochondrial mass. Likewise, the overall mitochondrial area per cell was not significantly affected (Fig. 3F). These results demonstrate that the increase in mtDNA copy number and the appearance of extramitochondrial DNA puncta in AltSLC35A4-KO cells occur independently of TFAM expression or mitochondrial mass, suggesting that AltSLC35A4 influences mtDNA organization through a distinct post-transcriptional, post-replication or structural mechanism rather than via the regulation of mtDNA transcription or replication.

**Figure 3:**
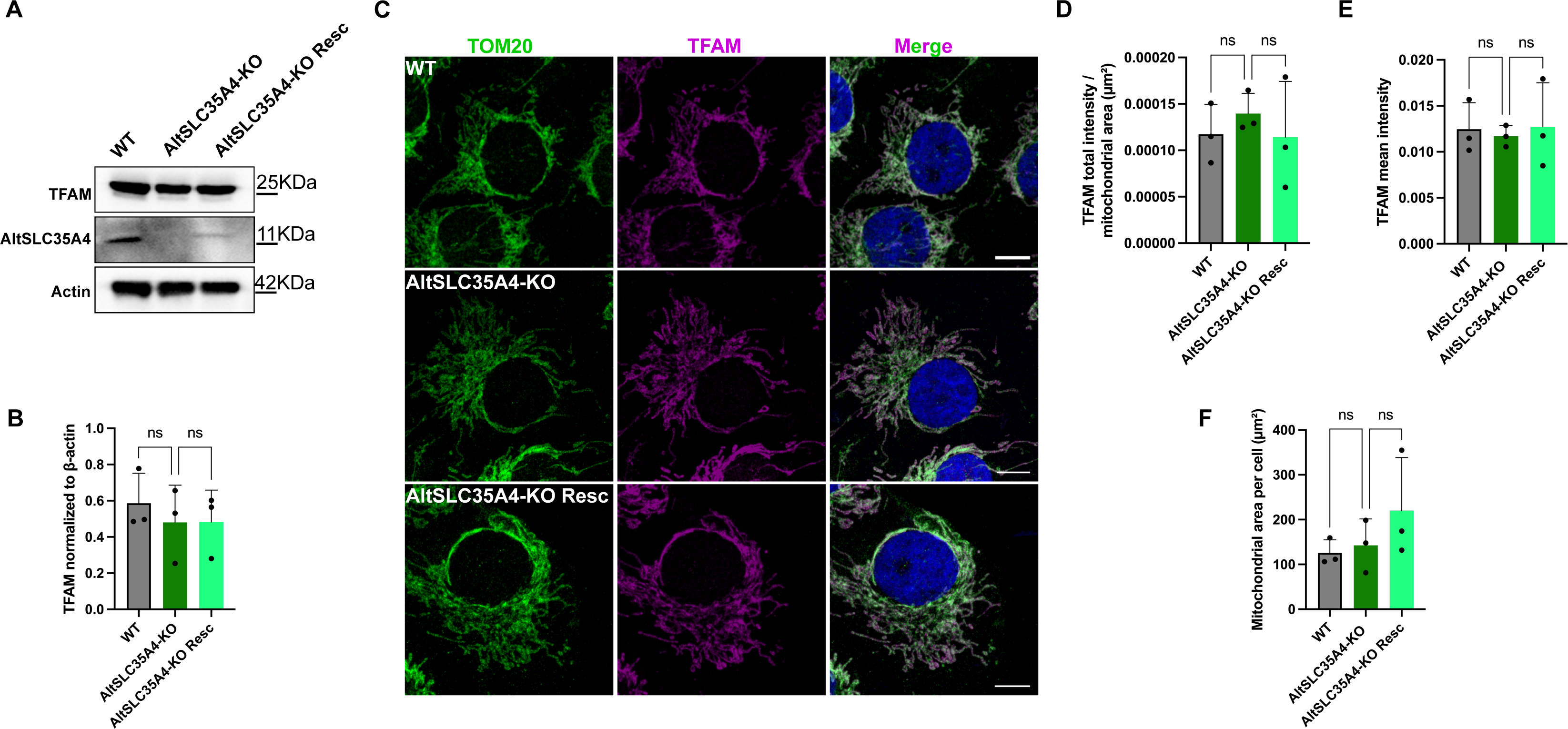
TFAM abundance and mitochondrial mass are unchanged in AltSLC35A4-deficient cells. (A) Representative immunoblot showing TFAM, AltSLC35A4, and β-actin levels in WT, AltSLC35A4-KO, and rescue cells. (B) Quantification of TFAM protein normalized to β-actin revealed no significant differences among conditions (mean ± SD, n = 3). (C) Representative confocal images of cells immunostained for TOM20 (mitochondria, green) and TFAM (magenta). Nuclei are counterstained with Hoechst (blue). Scale bars, 10 µm. (D–E) Quantification of TFAM fluorescence intensity normalized to mitochondrial area shows that both total and mean TFAM levels per mitochondrial area are comparable across genotypes. (F) The overall mitochondrial area per cell, measured from TOM20 signal, was not significantly different between groups. Data represent n=3 biological replicates. For image-based quantifications, between 78 and 203 cells were analyzed per replicate. Statistical significance was determined using one-way ANOVA; ns, not significant.

### Increased mtDNA copy number is not accompanied by changes in mitochondrial membrane potential

As mitochondrial membrane potential is a key determinant of organelle integrity and mtDNA retention and because alterations in mtDNA can also influence mitochondrial membrane potential ^21^, we next assessed whether the increase in mtDNA copy number and its release observed in AltSLC35A4-KO cells was associated with changes in mitochondrial membrane potential (Δψm). We measured JC-1 fluorescence in WT, AltSLC35A4-KO, and rescue cells. JC-1 is a potential-sensitive dye that accumulates in mitochondria in a voltage-dependent manner, forming red aggregates under high Δψm and emitting green fluorescence as monomers when the membrane potential decreases, while remaining confined to the mitochondrial network^22^. Confocal imaging and quantification of the JC-1 red (590 nm)/green (530 nm) fluorescence ratio revealed no significant differences among the three conditions (Fig. 4A–B), indicating that the mitochondrial network remains polarized despite the increased mtDNA copy number in the KO cells. Treatment of WT cells with the uncoupler FCCP, used as a positive control, caused the expected loss of JC-1 aggregates and a marked reduction of the 590/530 ratio (Fig. 4C–D). Together, these results indicate that variations in mtDNA copy number induced by AltSLC35A4 deficiency occur independently of changes in mitochondrial membrane potential, suggesting that the observed mtDNA dysregulation is not a secondary consequence of depolarization, nor causes defects in inner membrane potential.

**Figure 4:**
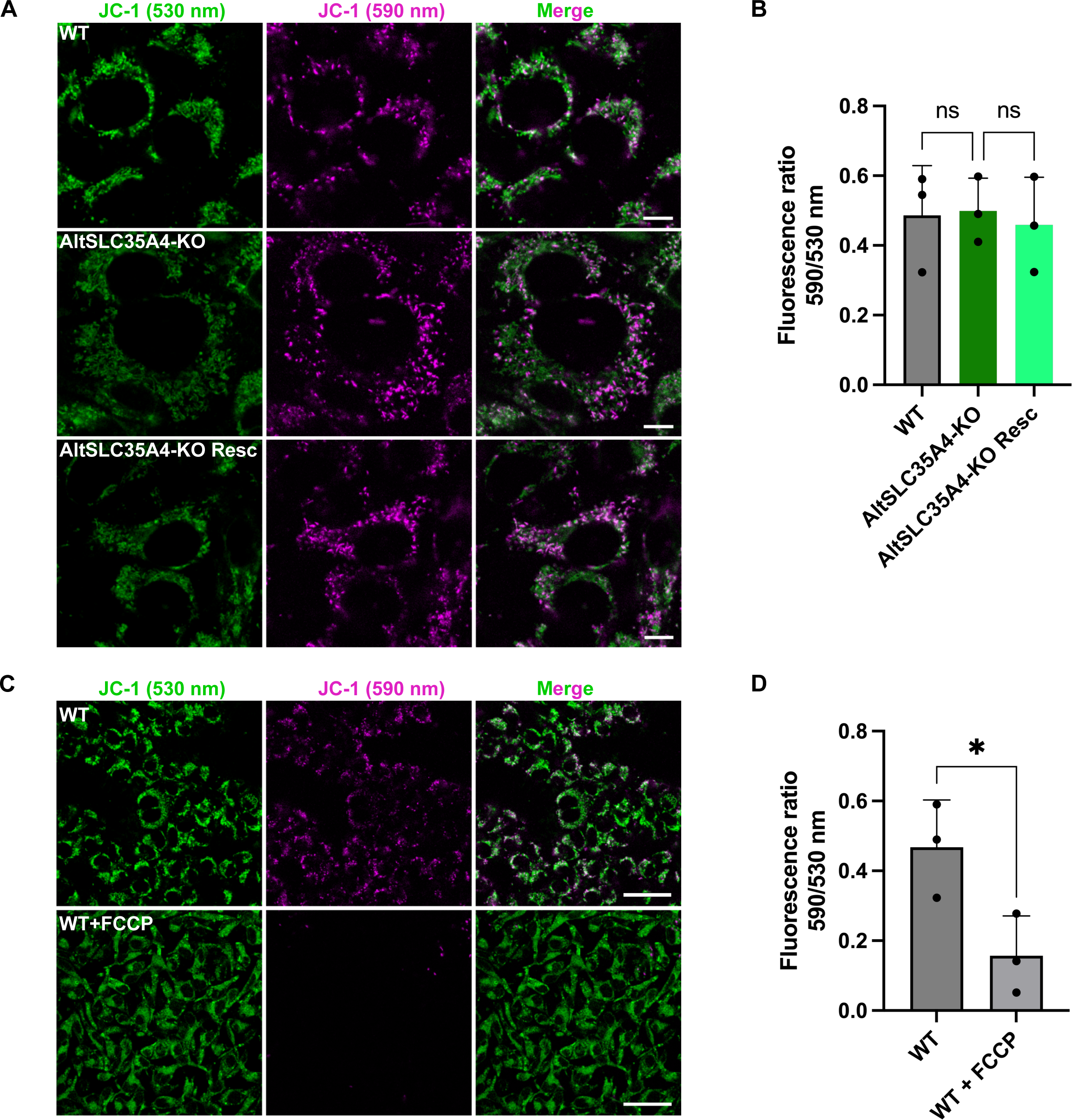
Assessment of mitochondrial membrane potential (Δψm) in AltSLC35A4-deficient cells. (A) Representative confocal images of WT, AltSLC35A4-KO, and rescue cells stained with the potential-sensitive dye JC-1. Green fluorescence (530 nm) corresponds to JC-1 monomers, while magenta fluorescence (590 nm) reflects J-aggregates formed at high Δψm. Scale bars, 10 µm. (B) Quantification of the red (590 nm)/green (530 nm) fluorescence ratio. (C) Validation of the assay using FCCP in WT cells, showing complete depolarization with loss of JC-1 aggregates. Scale bars, 100 µm. (D) Quantification of JC-1 ratios confirms the expected reduction upon FCCP treatment. Data represent mean ± SD from n = 3 independent experiments. Statistical significance was determined by one-way ANOVA; *, p < 0.05; ns, not significant.

### AltSLC35A4 deficiency triggers inflammatory signaling

Because the loss of AltSLC35A4 led to mtDNA accumulation and leakage, we next examined whether these mitochondrial alterations were accompanied by transcriptional signatures consistent with activation of the cGAS-STING pathway, which is known to be triggered by cytosolic mtDNA and induce downstream inflammatory transcriptional pathways^23^. We performed RNA sequencing on WT and AltSLC35A4-KO cells. The differential expression analysis identified 132 upregulated and 98 downregulated genes in AltSLC35A4-KO cells relative to WT (Fig. 5A). Gene ontology enrichment highlighted pathways involved in signal transduction, extracellular matrix organization, cell surface interactions and interleukin-4/interleukin-13 signaling (Fig. 5B), suggesting a broad activation of inflammation and immune-related transcriptional programs. To determine whether mitochondrial pathways were affected, we compared differentially expressed genes (DEGs) with the Mitocarta 3.0 list^24^. Only seven mitochondrial-associated genes overlapped with the differentially expressed set (Fig. 5C), indicating that mitochondrial transcriptional programs were largely preserved. Examination of normalized read counts for mitochondrial nuclear encoded genes confirmed that, AltSLC35A4-KO produced bidirectional changes, spanning mito-metabolic enzymes (*SQOR*, *ABAT*, *DGLUCY*, *CYP11A1*, and *MAOA)* and organelle dynamics/trafficking factors *(ARMCX2* and *SPIRE1)* (Fig. 5D). Heatmap analysis of mtDNA-encoded transcripts revealed a coherent upward trend in fold-change across the entire mitochondrial genome, consistent with the increased mtDNA copy number observed in AltSLC35A4-KO cells indicating that higher mitochondrial DNA abundance is paralleled by proportionally elevated mitochondrial RNA levels (Fig. 5E)

**Figure 5:**
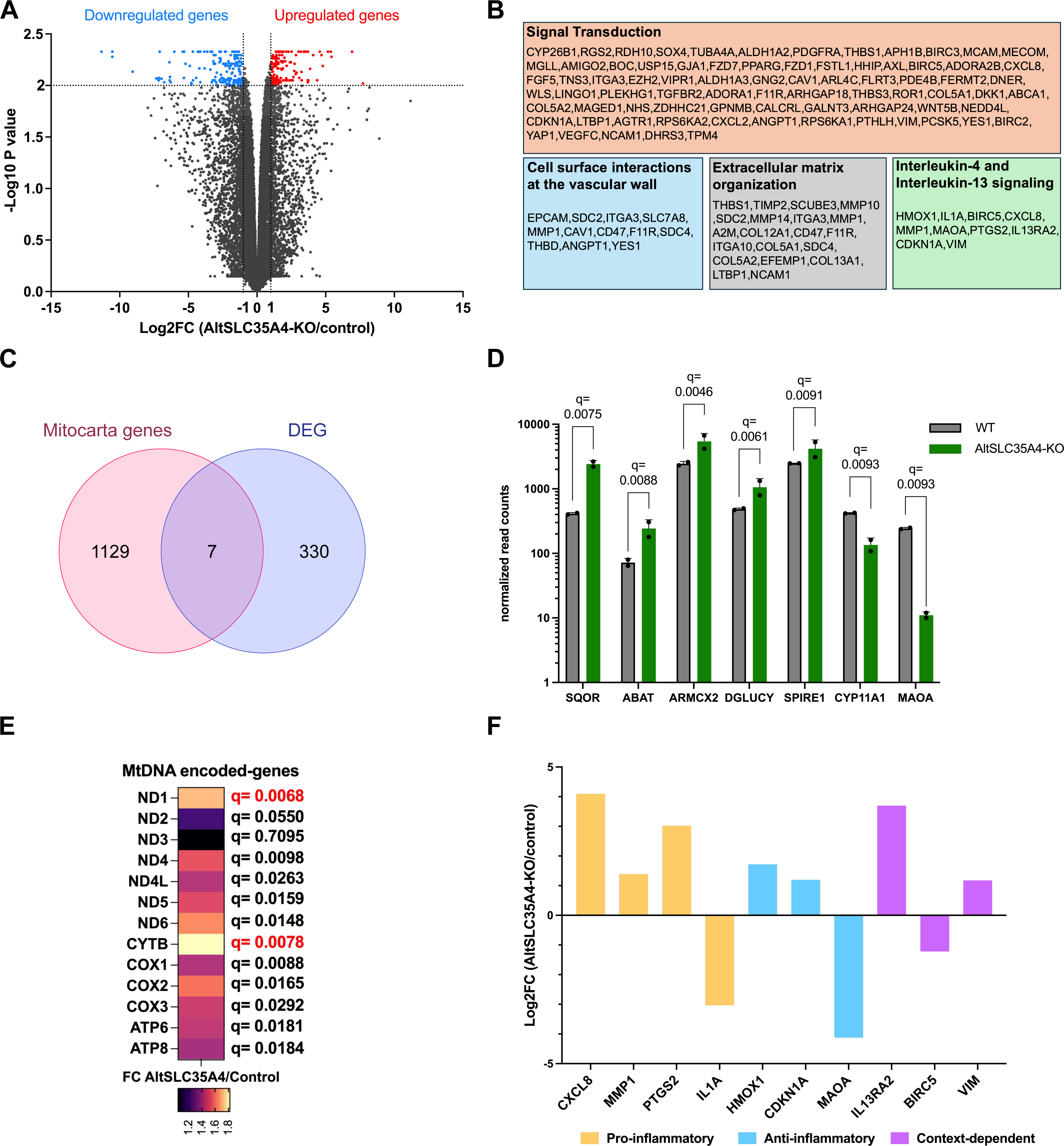
Loss of AltSLC35A4 rewires mitochondrial and inflammatory transcriptional programs. (A) Volcano plot showing differentially expressed genes (DEGs) between WT and AltSLC35A4-KO cells as detected by RNA-seq analysis. A total of 160 genes were upregulated and 177 downregulated (Log2FC ≥ 1, adjusted p < 0.01). (B) Gene Ontology enrichment analysis of DEGs highlights processes related to signal transduction, extracellular matrix organization, cell surface interactions at the vascular wall, and interleukin-4/interleukin-13 signaling pathways. (C) Venn diagram illustrating the overlap between Mitocarta 3.0 annotated mitochondrial genes and DEGs detected in AltSLC35A4-KO cells. (D) Normalized RNA-seq read counts for selected DEGs of nuclear-encoded mitochondrial genes. (E) Heatmap representing fold-change values for mitochondrial DNA encoded transcripts in AltSLC35A4-KO cells compared to WT, with associated q-values displayed for each gene, q-values in read represent genes that are DEG (F) Log2FC values for genes belonging to the IL-4/IL-13 signaling gene list. Genes are displayed according to their functional roles (pro-, anti-, or context-dependent inflammatory). Data are presented as mean ± SD. Q-values shown in (E) and (F) are derived from the RNA-seq differential expression analysis pipeline.

To further characterize the immune phenotype, we classified IL-4/IL-13 pathway genes according to their known functional categories (pro-inflammatory, anti-inflammatory, or context-dependent) (Fig. 5F). This revealed a mixed transcriptional profile, with the induction of pro-inflammatory mediators such as *CXCL8*, *MMP1*, and *PTGS2*, accompanied by regulatory or anti-inflammatory signals, including *HMOX1* and *CDKN1A*.

Together, these data suggest that loss of AltSLC35A4 triggers mtDNA-leakage–associated inflammatory remodeling alongside a compensatory bidirectional adjustments in nuclear mito-metabolic and organelle-dynamics programs pointing to an emerging mito-nuclear imbalance rather than a global failure of mitochondrial gene expression.

## Discussion

Our data identify AltSLC35A4 as a previously unrecognized component of mitochondrial genome homeostasis that connects the inner mitochondrial membrane to actomyosin machinery. Proteomics and imaging converge on MYH9/MYH10 as AltSLC35A4 partners, placing this alternative protein at the interface between mitochondria and actin networks. In this context, loss of AltSLC35A4 increases mtDNA copy number, raises nucleoid counts, and produces DNA puncta in extramitochondrial space, while leaving nucleoid size, TFAM abundance/localization, mitochondrial area, and Δψm unchanged. These features are mostly consistent with a maintenance defect, altered nucleoid organization and/or turnover rather than generalized mitochondrial dysfunction.

Mechanistically, we hypothesize that AltSLC35A4 contributes to positioning, segregation and remodeling of nucleoids via two non-exclusive models that fit the data. First, AltSLC35A4 could act as an IMM anchor or adaptor that couples nucleoids to actin/myosin forces required for their spacing and partitioning^1,18^. Loss of this linkage would promote an unscheduled replication (increase in mtDNA copy number) with occasional escape of nucleoids/DNA into the cytosol via a yet undefined mechanism. Alternatively, AltSLC35A4 could influence mtDNA degradation/turnover (e.g., exonuclease access or mitophagy routing). Impaired clearance would increase copy number and nucleoid counts without changing TFAM or Δψm, while leakage would explain the rise in extramitochondrial DNA puncta^25,26^. The selective nuclear transcriptional response we observe with enrichment for signal-transduction, extracellular matrix, and IL-4/IL-13 programs with an increase in mitochondrial transcripts supports a model in which mitochondrial perturbations are relayed to the nucleus via retrograde signaling without overt bioenergetic failure. Interestingly, oxidative stress is known to influence mtDNA release. For instance, ROS accumulation has been shown to trigger mtDNA leakage into the cytosol, activating downstream signaling pathways such as cGAS-STING and NF-κB that mediate inflammatory and stress responsive gene expression^27^. Considering that AltSLC35A4 protects against oxidative stress, the increased mtDNA copy number and extracellular mtDNA puncta observed in AltSLC35A4-KO cells may reflect subtle redox-dependent perturbations in mtDNA maintenance^15^.

These findings position AltSLC35A4 near known “mtDNA maintenance” nodes: (i) the actin DRP1/INF2 axis that shapes nucleoids and fission^6,28^; (ii) nucleoid-IMM tethers that govern spacing and^11,29^ (iii) pathways that define replication versus degradation set-points^30,31^. The lack of TFAM changes argues against a transcriptional/packaging cause and favors post-transcriptional/post-replication control. Notably, extramitochondrial mtDNA provide a plausible upstream source for innate immune activation, and together with the transcriptomic signature, they motivate direct testing of cGAS–STING activation, the central mediator of cytosolic DNA-induced innate immunity ^23,32^.

Alternative explanations remain possible. Increased copy number could reflect PGC-1α–driven biogenesis^33^ or decreased mitophagy^34^ rather than replication dysregulation *per se*. However, stable Δψm and mitochondrial area argue against a broad biogenic surge, especially since among Mitocarta 3.0 transcripts detected by RNA seq (=1110), 99.4% (=1103) were not differentially expressed and Mitocarta genes accounted for only 2.1% (=7) of all DEG. Together with unchanged TFAM levels and distribution, these observations make a broad transcriptional reprogramming unlikely. Likewise, the IL-4/IL-13 signature might arise from paracrine cues independent of mtDNA leakage^35^. Targeted experiments will discriminate among these possibilities.

We therefore propose the following testable predictions: (1) if cGAS STING pathway is activated, its pharmacologic or genetic inhibition (e.g., RU.521, H-151, cGAS/STING knockdown) will blunt the cytokine/extracellular matrix transcriptional response in AltSLC35A4-KO cells; (2) assays of mtDNA turnover (BrdU/EdU pulse-chase, mtDNA damage repair kinetics) will reveal altered renewal dynamics in KO cells consistent with perturbed replication-degradation balance; and (3) markers of selective mitophagy (PINK1/Parkin flux; mito-Keima) will distinguish turnover defects from replication-biased increases.

In sum, AltSLC35A4 emerges as a gatekeeper of mtDNA homeostasis that operates independently of TFAM and Δψm yet upstream nucleus-encoded stress programs. While our data support a conncection between AltSLC35A4, mitochondrial DNA maintenance and actomyosin-related pathways, the precise structural or mechanical basis of this regulation remains to be established. These findings nonethless open a mechanistic avenue for understanding how mitochondrial genome perturbations may interface with cellular immunity and tissue remodeling pathways.

## ACKNOWLEDGEMENTS

We thank the members of the Vanderperre lab for fruitful discussions and technical help. Many thanks to the technical experts from CERMO-FC technological platforms (Geneviève Bourret – Genomics; Farzaneh Rahmdani – Bioinformatics). We further thank Pr Gilles Gouspillou for providing the TFAM antibody.

Supplementary Table I.

This table provides a detailed list of primary and secondary antibodies used for Western blotting and immunofluorescence. For each antibody, the target antigen, commercial source, catalog number, host species, and application/dilution are indicated. Both HRP-conjugated and fluorescent secondary antibodies are included.

