## Supplementary Table I for "Regulation of mitochondrial DNA homeostasis by a mitochondrial microprotein"

### Supplementary Table 1

| Target | Company | cat # | species | application and dilution |
|---|---|---|---|---|
| **Primary antibodies** | | | | |
| HA Tag | Cell signaling | 3724S | Rabbit | IF (1/100), WB (1/1000) |
| HA Tag | Cell signaling | 2367S | Mouse | IF (1/100), WB (1/1000) |
| anti-DNA | Progen | 690014S | Mouse | IF (1/100), WB (1/1000) |
| TOM20 | Cell signaling | 42406S | Rabbit | IF (1/100), WB (1/1000) |
| TFAM | Abcam | ab119684 | Mouse | IF (1/100), WB (1/1000) |
| SDHA | Cell signaling | 5839S | Rabbit | WB (1/1000) |
| AltSLC35A4 | MediMabs (custom) | | Rat Polyclonal | WB (1/250) |
| beta-actin | Proteintech | 66009-1-Ig | Mouse | WB (1/10000) |
| Myosin IIA | Cell signaling | 49349S | Rabbit | WB (1/1000) |
| Myosin IIB | Cell signaling | 3404S | Rabbit | IF (1/100), WB (1/1000) |
| **Secondary antibodies (HRP)** | | | | |
| Anti-Rabbit HRP | Jackson ImmunoResearch | 712-035-153 | Donkey | WB (1/5000) |
| Anti-Mouse HRP | Jackson ImmunoResearch | 111-035-144 | Goat | WB (1/5000) |
| Anti-Rat HRP | Jackson ImmunoResearch | 115-035-146 | Goat | 000) |
| **Secondary antibodies (fluorescent)** | | | | |
| Anti-Rabbit Alexa 594 | Abcam | ab150116 | Goat | IF (1/1000) |
| Anti-Rabbit Alexa 488 | Jackson ImmunoResearch | 111-545-003 | Goat | IF (1/1000) |
| Anti-Mouse Alexa 647 | Abcam | ab150116 | Goat | IF (1/1000) |

IF=immunofluorescence
WB=Western blotting
